# Protein Biomarker in Focal Cortical Dysplasia: Molecular Clues to Pathogenesis

**DOI:** 10.1101/2025.09.27.678943

**Authors:** Priyanka Soudarpally, Syed Sultan Beevi, Vinod Kumar Verma, Anuja Patil, Manas Panigrahi, Madigubba Sailaja, Radhika Chowdary Darapuneni, Tincy A John, Sita Jayalakshmi

**Affiliations:** Department of Neurology, Krishna Institute of Medical Sciences, Minister Road, Secunderabad, Telangana, India; Diagnostics Division, Krishna Institute of Medical Sciences, Minister Road, Secunderabad, Telangana, India; KIMS Foundation and Research Centre, KIMS Hospital, Minister Road, Secunderabad, Telangana, India

**Keywords:** FCD, drug-resistant epilepsy, mTOR, AKT, DEPDC5, KCNT1, precision medicine

## Abstract

Focal Cortical Dysplasia (FCD) is a major cause of drug-resistant epilepsy (DRE), particularly in pediatric and young adult populations characterized by structural abnormalities in cortical development. This study investigated 60 patients with histologically confirmed FCD, combining clinical, histopathological, and molecular data to identify subtype-specific molecular signatures. we investigated a panel of candidate protein biomarkers (AKT, PTEN, mTOR, HTR6, RHEB, KCNT1, RALA, DEPDC5) across FCD subtypes using patient-derived tissue samples.

Our results reveal subtype-specific alterations in biomarker expression, particularly within the mTOR signaling pathway, supporting its central role in cortical malformations.

Key dysregulated genes AKT, mTOR, PTEN, RHEB, DEPDC5, KCNT1, RALA, and HTR6 involved in mTOR signaling, neuronal excitability, and cortical development. Western blotting and IHC revealed marked upregulation of AKT and mTOR in FCD III, consistent with mTOR pathway hyperactivation.

In contrast, KCNT1, RALA, and DEPDC5 were significantly downregulated across all subtypes, suggesting disrupted inhibitory signaling and GATOR1 complex dysfunction. ELISA assays validated increased expression of AKT, mTOR, and HTR6, particularly in higher-grade lesions. This study bridges clinical, histopathological, and protein-level data, providing novel insight into the molecular basis of FCD and highlighting candidate biomarkers for future diagnostic and therapeutic applications

## 1. Introduction

Focal cortical dysplasia (FCD) is a heterogeneous malformation of cortical development and one of the leading causes of drug-resistant epilepsy in both children and adults. Histopathological classification according to the International League Against Epilepsy (ILAE, 2011) defines subtypes based on cortical disorganization, dyslamination, and cellular abnormalities such as balloon cells. Epidemiological data indicate that FCDs account for approximately 5%–25% of epilepsy cases globally, affecting an estimated 50 million individuals worldwide (1). Clinically, FCD presents with a spectrum of manifestations, including drug resistant seizures, cognitive delays, developmental regression, and behavioral disturbances, leading to substantial morbidity and psychosocial burden (2).

The etiology of FCD remains multifactorial such as prenatal infections or trauma, suggests a crucial role for somatic and germline mutations in key regulatory genes (3). Previous studies have identified genomic variants in mTOR pathway genes (e.g., MTOR, AKT3, RHEB, DEPDC5, TSC1, TSC2) and other loci implicated in neuronal signaling, ion channels, and neurodevelopment (4). Hyperactivation of the mammalian target of rapamycin (mTOR) signaling pathway has emerged as a fundamental mechanism in the pathogenesis of FCD. MTOR regulates crucial cellular processes like growth, differentiation, autophagy, and synaptogenesis (5).

Despite the use of advanced neuroimaging modalities such as high-resolution MRI, FDG-PET, and SPECT to localize epileptogenic lesions, many cases of FCD, especially those classified as MRI-negative, remain difficult to diagnose with certainty (6). These limitations underscore the pressing need for candidate molecular biomarkers that can enhance diagnostic precision, stratify FCD subtypes, and guide clinical management. While several studies have investigated transcriptomic and proteomic changes in FCD tissue, a consensus on reliable, reproducible biomarkers is lacking (7). In addition to the canonical mTOR pathway genes, newer studies have identified additional candidates such as *IRS1, HTR6, ZNF337*, and *RAB6B*, which may influence mTOR signaling and play a role in cortical dysgenesis. These findings suggest that a network-level approach involving multiple genes may be necessary to capture the molecular heterogeneity of FCD (8).

A significant lacuna in the field is the lack of comparative molecular profiling across different FCD subtypes, which limits our understanding of subtype-specific pathophysiology (8). Another gap lies in the lack of validation using standardized bioinformatics pipelines that integrate genomic, transcriptomic, and proteomic datasets. Thus, there is a critical need to identify biomarkers that not only correlate with FCD but are also subtype-specific and reproducible across cohorts.

The present study aims to address these gaps by identifying a panel of molecular biomarkers with diagnostic and pathophysiological relevance to FCD. Based on comprehensive literature and in silico screening, eight candidate genes have been shortlisted: *AKT, PTEN, mTOR, HTR6, RHEB, KCNT1, RALA*, and *DEPDC5* (9). These genes were selected through integrative analysis using multiple bioinformatics tools and databases, including the UCSC Genome Browser, GTEx, GeneCards, DAVID, STRING, Enrichr, ClinVar, gnomAD, PolyPhen, SIFT, MutationTaster, dbSNP, ExAC, VarSome, HGMD, and Ensembl (10). The selection criteria included known involvement in FCD, seizure disorders, mTOR pathway regulation, and expression overlaps with renal angiomyolipoma lesion that also demonstrates aberrant mTOR signaling.

The proposed study will involve collecting and analysis of brain tissues from histologically confirmed cases of FCD Types I, II, and III. Protein expression of the selected markers will be assessed using immunoblotting and immunohistochemistry to determine spatial and quantitative differences across FCD subtypes. This approach will allow identification of both common and subtype-specific markers. Furthermore, gene interaction and network analysis will be performed to elucidate potential co-regulatory mechanisms and signaling crosstalk, providing deeper insight into disease pathogenesis. Additionally, to situate our findings within the broader molecular landscape, we reviewed published genomic studies and compiled literature-derived variants and signaling pathways previously implicated in FCD. By clearly distinguishing between patient-derived biomarker data and literature-derived contextual information, this study provides a comprehensive and translational framework for understanding molecular signatures of FCD. Ultimately, the research aims to bridge the current gap between clinical diagnosis and molecular pathology in FCD, contributing to a more personalized and mechanistic approach to epilepsy management.

## 2. Materials and Methods

### 2.1 Patient cohort description and sample collection

We analyzed resected cortical tissue from 60 patients diagnosed with FCD and undergoing epilepsy surgery at KIMS Hospitals, Secunderabad, with due informed consent. The cohort included 34 males and 26 females, with ages ranging from 6 to 42 years (mean ± SD: 21.4 ± 8.6 years) (supple. Table 1). Seizure types included focal impaired-awareness seizures (68%), focal-to-bilateral tonic-clonic seizures (22%), and mixed phenotypes (10%). Localization of epileptogenic zones was predominantly frontal (55%), temporal (30%), and parietal/occipital (15%). Histological classification was performed according to ILAE 2011 criteria: FCD type I (n = 18), type II (n = 28), and type III (n = 14) (Supple. Table 2).

Freshly resected samples were collected in carrier media, processed and stored in RNA later at −80 °C. All the samples were collected with prior approval from the Ethics Committee of Krishna Institute of Medical Sciences.

### 2.2 Immunohistochemistry

Two µm-thick FFPE sections were cut for each block, deparaffinized in xylene, and rehydrated through graded ethanol and water. Sections were equilibrated in PBS and subjected to endogenous peroxidase blocking with 1.5% H□O□ for 45 minutes. Antigen retrieval was performed by boiling sections in 10 mM Tris-EDTA (pH 9.0) for 20 minutes. The sections were blocked with 4% bovine serum albumin (BSA) in PBS for 60 minutes and incubated overnight at 4 °C with primary antibodies diluted in the blocking solution. Primary antibodies included AKT3, HTR6, PTEN, DEPDC5 (rabbit polyclonal, 1:100 dilution) and MTOR, PTEN, KCNT1, RALA, RHEB (mouse monoclonal, 1:100 dilution) were used for incubation O/N at 4°C. After washing with PBS, slides were incubated with HRP-conjugated secondary antibodies at room temperature for 1 hour. The slides were then exposed to HRP substrate DAB, counterstained with hematoxylin, and mounted using DPX. Slides were then visualized in bright field mode at 63X magnification using an Olympus microscope (OLYMPUS BX53). Differential staining was assessed manually in at least four microscopic fields, with scores averaged for target protein expression.

### 2.3 Western Blotting

Protein lysates from fresh tissue samples were extracted using RIPA lysis buffer with 1% phenylmethylsulphonyl fluoride (PMSF), 1% protease inhibitor and incubated for 30 minutes on ice. Followed by ultrasonication for 2 minutes on ice with an interval after every 5 sec and repeated once. Then incubate on ice for 15 minutes. Centrifuged at 14000 g for 20 min at 4□. Supernatant was collected, used to determine the protein concentration using Follin-Lowry method. Tissue lysates of 25µg +1x sample buffer are boiled at 95□ and resolved via SDS-PAGE using 8%,10%, w/v gels following electrophoresis, proteins were transferred onto PVDF membranes for 90 min at 150mv, then blocked with 5% BSA in PBST for 1 hour at RT to prevent non-specific binding. The membranes were subsequently incubated overnight at 4□°C with a panel of primary antibodies, including rabbit polyclonal antibodies against 5-HT6R, AKT3, PTEN, and DEPDC5 (all at 1:1000 dilution), and mouse monoclonal antibodies against MTOR, KCNT1, RALA, and RHEB (also at 1:1000). β-Actin (1:5000) was used as an internal loading control. After thorough washing with PBST, membranes were incubated with HRP-conjugated secondary antibodies (anti-rabbit or anti-mouse, as appropriate) at a dilution of 1:5000 for 1 hour at RT. Chemiluminescence was detected using the Electrochemiluminescence (ECL) kit (Bio-Rad, Clarity Max ECL Substrate) and a Syngene electrophoresis image analyzer. Densitometric analysis was done using Image J software and comparative density plots were represented as Bar diagrams.

### 2.4 Tissue Controls

Given the high expression of mTOR in proliferative germ cells with active mTOR signaling pathways, human testis tissue was utilized as positive control for mTOR and DEPDC5. KCNT1 antibody specificity was confirmed using human prostate cancer tissue, which has been reported to express voltage-gated potassium channels at elevated levels. For HTR6, human brain tissue was used, reflecting its physiological expression in central nervous system regions associated with serotonin-mediated signaling. RALA expression was verified in human ovarian cancer tissue, where Ras-related GTPases are commonly dysregulated. PTEN was validated in human breast cancer tissue, known for altered PI3K/AKT signaling and frequent PTEN mutations or loss. AKT3 in mouse brain and RHEB in rat brain, both of which are highly enriched in the cerebral cortex and are integral to mTOR pathway regulation in neuronal development and function. These control tissues ensured accurate antigen localization and confirmed antibody reliability for downstream comparative analysis in FCD patient samples.

### 2.5 ELISA

Direct ELISA was tissue lysates of type I, II & III FCD samples as antigen and primary antibodies against HTR6, AKT, KCNT1, and mTOR, begin by coating a 96-well ELISA plate with tissue lysate (diluted appropriately in carbonate-bicarbonate buffer, pH 9.6) and incubated overnight at 4°C. After washing PBS Tween (0.05% Tween-20), blocked non-specific binding sites with 5% BSA in PBS for 1 hour at RT. Following blocking, incubated the antigen coated wells with HRP-conjugated primary antibodies specific for HTR6, AKT, KCNT1, and mTOR, diluted in blocking buffer to the wells in triplicates for each primary antibody and incubated for 1–2 hours at RT. Washed the wells thoroughly to remove unbound antibodies, then added TMB substrate and incubated in the dark until color develops. Reaction was stopped using 1N HCl or 2N H□SO□ and measured absorbance at 450 nm using a microplate reader (Agilent Technologies). Secondary antibody controls and blocking buffer controls were used to normalize the data.

### 2.6 Bioinformatics analysis

To investigate the biological significance and potential interactions among genes associated with focal cortical dysplasia (FCD) a comprehensive bioinformatics approach was used. Eight genes (MTOR, RALA, RHEB, DEPDC5, KCNT1, AKT3, HTR6, and PTEN) known to be associated in various FCD subtypes were selected based on prior literature, experimental and clinical evidence. Protein-protein interaction (PPI) analysis was performed using the STRING v12.0 database. Initially, interactions were assessed at the default medium confidence level (score # 0.4), but limited connectivity was observed due to sparse curated or experimental data. To capture a more comprehensive network, the confidence score was lowered to 0.15, and additional interactors were incorporated using the STRING “+ more” function, enhancing network complexity and enabling downstream enrichment analysis. Functional pathway enrichment was conducted using ShinyGO v0.80, analyzing the input genes across KEGG, Gene Ontology: Biological Processes (GO-BP), and Molecular Function (GO-MF).

### 2.7 Statistical analysis

The results are displayed as percentage of mean ± standard deviation (SD), as specified in the figure legends. Data analysis involved the use of appropriate statistical methods, to calculate the significance level and p value. All statistical analyses were with a significance level set at p<0.05, indicating statistical significance.

## 3. Results

### 3.1 Selection of FCD candidate markers using bioinformatics tools and databases

The selection of genes involved in focal cortical dysplasia (FCD) was made using an in-silico approach to identify genes with mutations that are functionally linked to the pathophysiology of FCD. By utilizing databases like PubMed, NCBI Gene, and PMC, along with gene pathway analysis, we identified key genes involved in the mTOR pathway (such as MTOR, RHEB, DEPDC5, PTEN, and AKT3), which play significant roles in neuronal development, cortical malformations, and epilepsy. Additionally, genes like KCNT1, involved in ion channel regulation, and RALA, part of the RAS signaling pathway, were included for their potential contributions to neuronal excitability and development, even though their direct roles in FCD are less well-defined (Supple. table 3). The genes were selected based on evidence of somatic mutations or pathway dysregulation contributing to FCD in various clinical and genetic studies, highlighting their functional relevance in both the molecular mechanisms of cortical dysplasia.

### 3.3 Expression of FCD candidate markers

Western blotting and immunohistochemistry (IHC) methods were used to investigate the differential expression of eight key molecular markers such as AKT, PTEN, KCNT1, RALA, mTOR, HTR6, RHEB, and DEPDC5 across histologically qualified focal cortical dysplasia (FCD) subtypes (Types I, II, and III) in comparison with positive and negative controls. The analyses revealed a significant upregulation of AKT and MTOR in FCD Type III lesions, demonstrated by strong protein band intensities in western blots and peculiar cytoplasmic and nuclear staining in IHC, particularly within dysmorphic neurons and balloon cells. Densitometric quantification corroborated these findings, with statistically significant increases (p < 0.05) in their expression along with IHC scoring in Type III relative to types I and II (Fig 1A and supple. table 4). These data suggest hyperactivation of the PI3K-AKT-mTOR pathway, which is known to promote aberrant neuronal growth, increased cell size, and epileptogenic, typical features of advanced FCD. In contrast to previous literature reporting loss of PTEN function, our study revealed a moderate level of upregulation in FCD Types II and III, as compared to Type I and positive controls, in both western blotting and IHC (Fig.1A and supple. table 4). This apparent contradiction may indicate compensatory feedback mechanisms attempting to reflect complex subtype-specific regulation of PTEN post-translational activity rather than total expression levels. HTR6, a G protein-coupled serotonin receptor involved in neurodevelopment and cortical excitability, exhibited mild expression level across all FCD subtypes as compared to positive control both in western blotting and IHC (Fig.1A and supple. table 4). This pattern suggests that HTR6 may be more relevant in physiological rather than dysplastic cortical signaling and impaired neurodevelopmental in FCD lesions. RHEB, a small GTPase that directly activates mTORC1, was minimally expressed in type II & III samples, further reinforcing the centrality of mTOR hyperactivity in advanced FCD pathology (Fig.1A and supple. table 4). Whereas the expression of RALA, a Ras family GTPase implicated in vesicular trafficking and synaptic function, and KCNT1, a sodium-activated potassium channel gene associated with neuronal excitability and epilepsy, was consistently downregulated across all FCD types, suggesting a loss of regulatory control over excitatory signaling in dysplastic cortex (Fig.1B and supple. table 4). Similarly, DEPDC5, a core component of the GATOR1 complex that negatively regulates mTORC1 in response to amino acid signaling, was markedly downregulated in all FCD subtypes (Fig.1B and supple. table 4). This observation shows that DEPDC5, a negative regulator of mTORC1, aligns with the hypothesis of mTOR hyperactivation in FCD, while reduced KCNT1 and RALA expression may contribute to abnormal neuronal excitability and impaired signal transduction. Densitometric quantification further validated these observations, demonstrating significantly reduced expression levels (p < 0.05) of KCNT1, RALA, and DEPDC5 in FCD lesions, independent of subtype.

**Figure 1.**
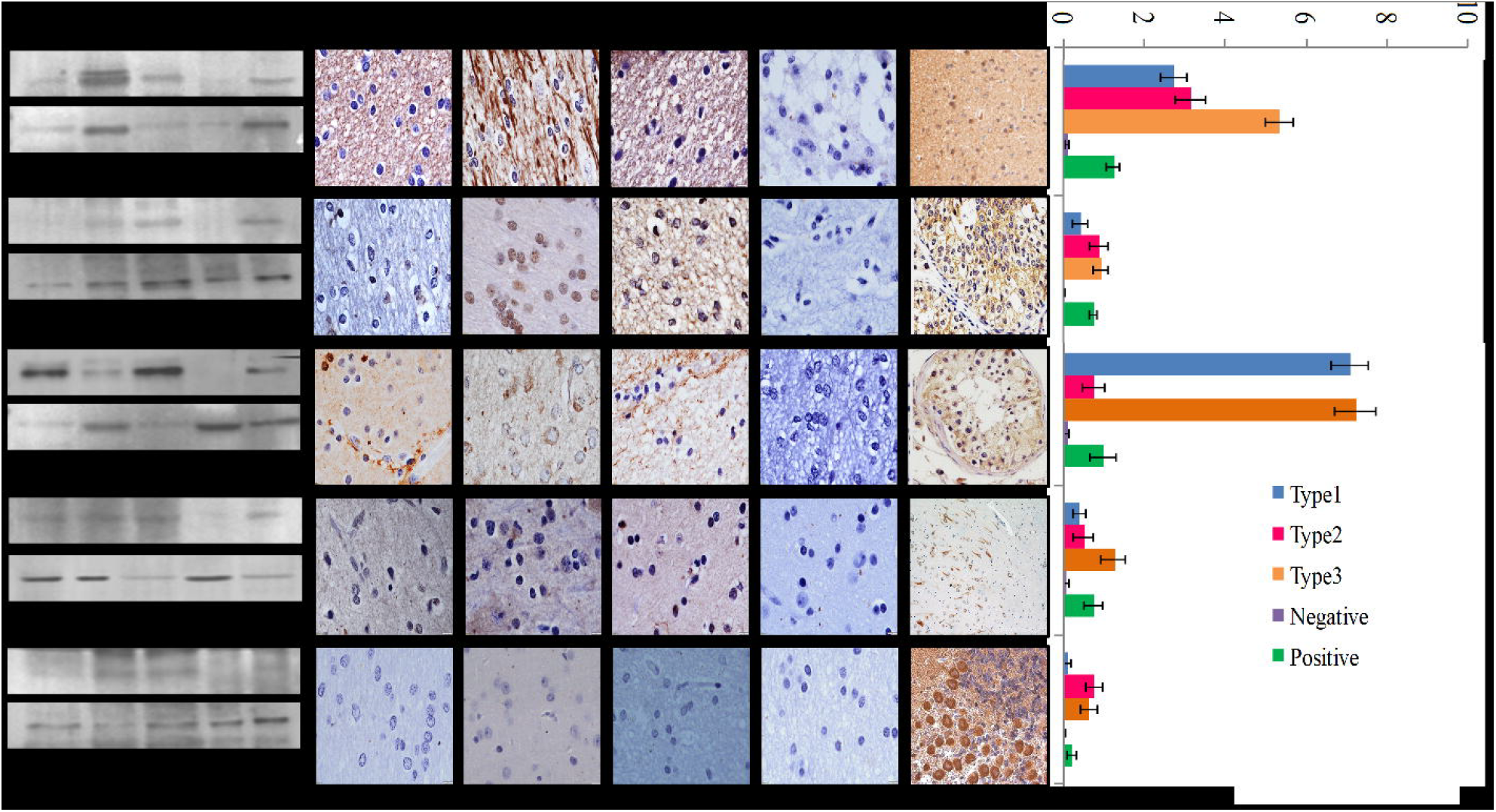
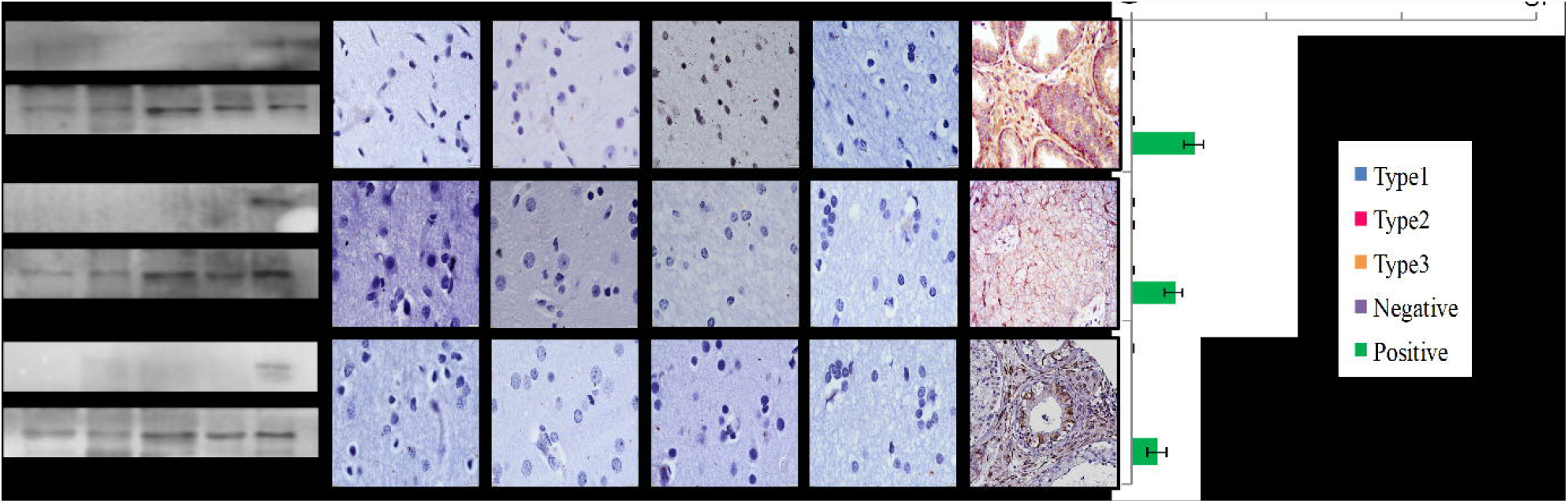
**A& B:** Representative Western blot, immunohistochemistry (IHC), and densitometric analyses of AKT, PTEN, KCNT1, RALA, MTOR, HTR6, RHEB, and DEPDC5 expression across focal cortical dysplasia (FCD) subtypes (Type I, II, III), negative controls, and positive controls. Western blot panels (left) show differential protein expression patterns, while corresponding IHC images (center) display cellular localization and intensity of immunoreactivity in cortical tissue sections. Quantitative densitometry analysis (right bar graph) represents relative expression levels ± SE for each marker across groups. All the experiments were repeated three times. Scale bar for IHC images: 50 µm.

### 3.4 Validation of positive FCD markers

Enzyme-linked immunosorbent assay (ELISA) was performed to validate the expression levels of selected FCD positive markers like AKT, mTOR, and HTR6 across FCD subtypes. The results demonstrated that mTOR levels were significantly elevated in FCD Type 3 compared to Types 1 and 2, with Type 1 also showing moderate expression. AKT concentrations increased progressively from Type 1 to Type 3, supporting its upregulation in more severe subtypes. HTR6 levels showed a modest but consistent rise across the subtypes, with the highest expression in Type 3. These ELISA findings verify the findings of immunoblot and immunohistochemistry confirming AKT, mTOR, and HTR6 as positive markers with subtype-specific differential expression in FCD pathology (Fig.2).

**Figure 2:**
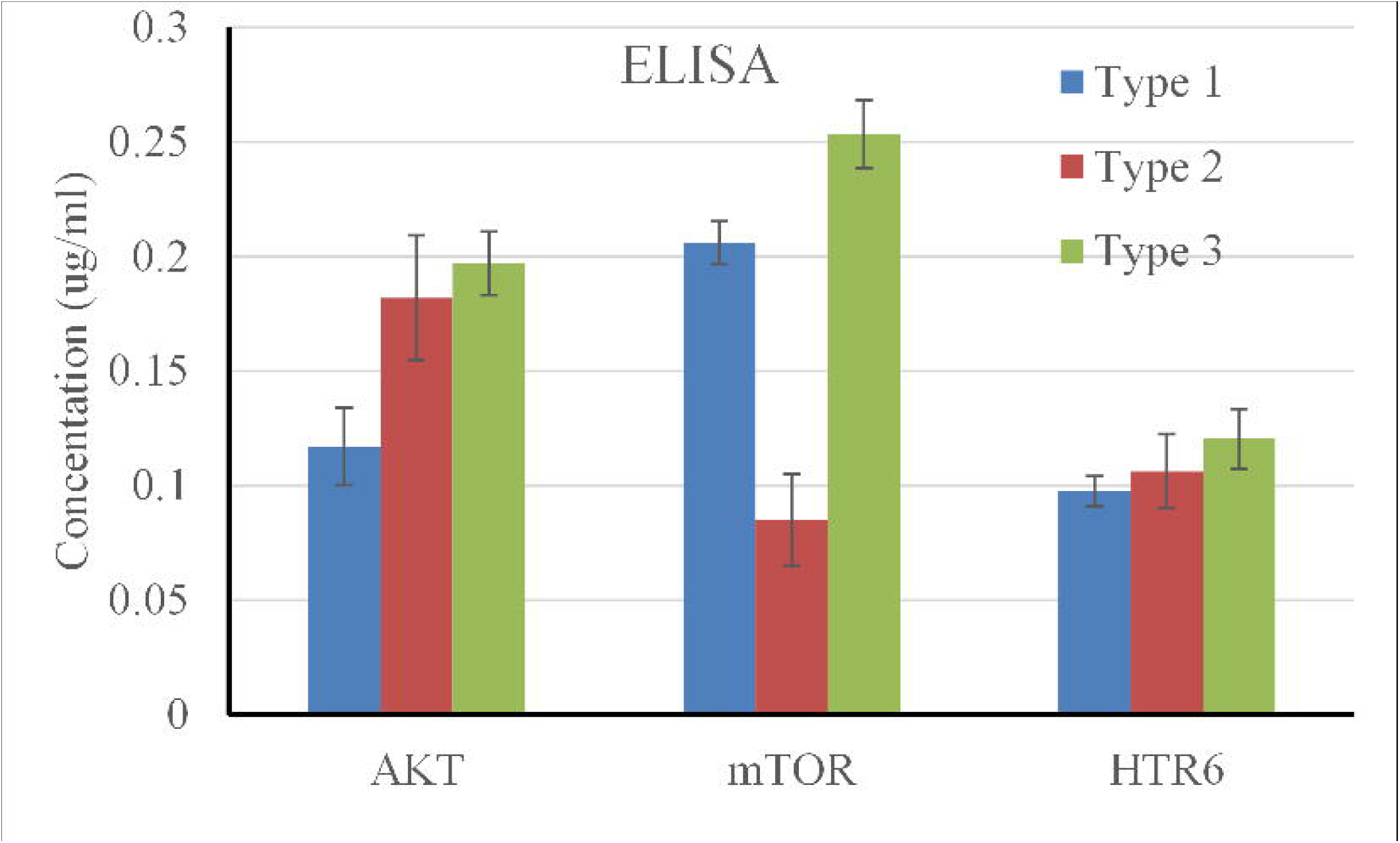
Differential expression of AKT, mTOR, and HTR6 proteins in FCD subtypes as measured by ELISA. *P*rotein concentrations (µg/mL) of AKT, mTOR, and HTR6 were quantified in tissue samples from patients with FCD Type I (blue), Type II (red), and Type III (green) using enzyme-linked immunosorbent assay (ELISA). Data represents standard deviation from biological replicates.

## 4. Discussion

This study identifies candidate protein biomarkers with distinct expression patterns across FCD subtypes, most prominently implicating mTOR signaling. Our findings are consistent with prior genomic studies reporting pathogenic variants in MTOR, AKT3, RHEB, DEPDC5, TSC1, and TSC2, further linking protein-level alterations with established genetic mechanisms. Molecular profiling highlighted aberrations in the PI3K-AKT-MTOR signaling pathway, a key regulator of cell proliferation, metabolism, and synaptic plasticity (11). Increased expression of AKT, MTOR, and RHEB was observed in FCD Type III, indicating pathway hyperactivation (12). These findings are consistent with previous studies that describe mTOR hyperactivity as a hallmark of malformations of cortical development (13). Interestingly, PTEN, a tumor suppressor and negative regulator of mTOR, showed moderate upregulation in FCD II and III, possibly as a feedback response to upstream mTOR activation (14). Gene expression analysis also revealed downregulation of DEPDC5, a component of the GATOR1 complex that suppresses mTORC1 activity (15). DEPDC5 mutations have been strongly associated with familial focal epilepsy and sporadic FCD, especially when “second hit” somatic mutations are present (16). Our data showed marked downregulation of DEPDC5 in all FCD subtypes, supporting its role as a critical gatekeeper in mTOR regulation.

Other dysregulated genes included KCNT1, a sodium-activated potassium channel linked to autosomal dominant nocturnal frontal lobe epilepsy and malignant migrating partial seizures of infancy (17). Its consistent downregulation across all FCD types may contribute to the hyperexcitability of neurons. Similarly, RALA, involved in synaptic vesicle trafficking, was also downregulated, implicating impaired neurotransmission in FCD pathophysiology (18). Gene interaction network analysis using STRING and GeneMANIA tools revealed that these genes such as MTOR, AKT3, DEPDC5, PTEN, RHEB, and KCNT1 form a tightly regulated network converging on the mTOR signaling axis (19). Disruption in any component can lead to a cascade of dysregulated neurodevelopmental processes, culminating in abnormal cortical lamination, dysmorphic neurons, and balloon cells, as observed in FCD Type IIb.

Clinically, these molecular changes correlate with seizure severity and lesion type. Patients with FCD Type III (often associated with hippocampal sclerosis or tumors) had more complex genetic alterations, longer seizure duration, and required surgical resection for seizure control (20). On the other hand, FCD Type I patients had subtler molecular changes and more focal epileptogenic zones (21). Disease associations from literature support the involvement of these genes not only in FCD but also in other neurological disorders such as tuberous sclerosis complex (TSC), megalencephalic, and hemimegaloencephaly. Notably, mTOR and RHEB mutations are found in both FCD and TSC, further emphasizing a shared molecular etiology (22). A thorough understanding of gene networks and patient-specific mutations will be key to developing precision therapies, including the use of mTOR inhibitors like rapamycin. Importantly, this study provides direct experimental evidence of biomarker dysregulation in surgically resected patient tissue, whereas the genomic variants and signaling pathways are literature-derived and included for contextual comparison only.

Our findings highlight translational potential subtype-specific candidate protein biomarkers which may support diagnostic classification and targeted interventions, particularly as mTOR inhibitors are under clinical investigation for related epileptic encephalopathies. Future functional studies should validate causality through pathway inhibition, gene silencing, or CRISPR-based correction models.

## 5. Conclusion

This study identifies subtype-specific dysregulation of candidate protein biomarkers in FCD, particularly highlighting mTOR hyperactivation in FCD II. These findings are derived directly from patient tissue and add a crucial protein-level dimension to the understanding of cortical dysplasia. By integrating patient-derived protein evidence with literature-derived genomic and pathway data we provide a comprehensive framework linking histopathology, molecular mechanisms, and potential therapeutic targets. Future studies should pursue functional validation and integrate genomic and proteomic analyses for improved biomarker-driven classification of FCD.

## Supporting information

legned

## Acknowledgement

We would like to express our gratitude to KIMS Hospitals Neurology team for their assistance in gathering patient information and samples.

## Approval

Ethics Committee, KIMS Hospitals (KIMS/ECBMHR/2022/33-01A), Minister Road, Secunderabad, Hyderabad, Telangana, India.

## Authors Contribution

The conceptualization, experimental design and data analysis was conducted by VKV and SSB, who also authored the original draft. SJ, MP and AP conceptualization, patient screening, information, sample collections and finalization of original draft. PS execution and data collection and compilation. TAJ performed gene mining and variant and data analysis.MS & RC patient stratification, facilitation of histological studies and finalization of manuscript.

## Supplementary table legend

**Table 1:** Distribution of individuals across different age groups by gender, showing female and male counts, total number of individuals, and corresponding percentage of the total population (N = 60).

**Table 2:** This table summarizes key clinical and demographic features associated with various types of Focal Cortical Dysplasia (FCD). It includes FCD subtypes (e.g., FCD I, II, IIIa, MTS combinations) along with their common clinical features such as focal motor seizures, behavioral arrest, automatisms, and generalized tonic-clonic seizures. The seizure types vary across subtypes, including complex partial seizures, ictal smiling, and focal to bilateral tonic-clonic seizures. The age range of affected patients spans from early childhood to late adulthood (3–58 years). The table also provides the approximate percentage of patients exhibiting each FCD subtype, reflecting their relative prevalence within the studied cohort.

**Table 3:** This table lists key genes associated with Focal Cortical Dysplasia (FCD) and related seizure disorders. It includes gene names, associated syndromes or conditions, and expression or frequency data. Each gene’s selection is justified based on its known role in neuronal development, mTOR signaling pathway regulation, or contribution to seizure susceptibility and cortical malformations. References are cited to support the gene-disease associations and the rationale for their inclusion in FCD-related genetic studies.

**Table 4:** This table presents the quantitative expression levels (mean ± standard deviation) of selected molecular markers across different subtypes of Focal Cortical Dysplasia (FCD I, II, and III). The markers include genes involved in the mTOR signaling pathway (AKT3, PTEN, MTOR, RHEB), serotonin signaling (HTR6), ion channel regulation (KCNT1), and other epilepsy-related genes (RALA, DEPDC5). The values indicate differential gene expression, suggesting subtype-specific molecular signatures.

