## Supplementary material for "Protein Biomarker in Focal Cortical Dysplasia: Molecular Clues to Pathogenesis": table

**Supplementary table 1: FCD patients’ distribution**

| **Age Group (years)** | **Female Count** | **Male Count** | **Total** | **Percentage (%)** |
| --- | --- | --- | --- | --- |
| **0–10** | 3 | 6 | 9 | 15.00 |
| **11–20** | 9 | 9 | 18 | 30.00 |
| **21–30** | 7 | 8 | 15 | 25.00 |
| **31–40** | 4 | 6 | 10 | 16.67 |
| **41+** | 2 | 6 | 8 | 13.33 |
| **Total** | 25 | 35 | 60 | 100.00 |

**Supplementary table 2: FCD patient types, clinical history and incidence**

| **FCD Type** | **Common Clinical Features** | **Seizure Types** | **Age Range (Years)** | **% of patients** |
| --- | --- | --- | --- | --- |
| **FCD I + II** | Refractory epilepsy, focal motor seizures, hyperactivity, behavioral arrest | Focal Motor (±BTCS), Hypomotor, Automotor, Complex Partial, Vocalization | 3–58 | ~60% |
| **FCD IIb** | Jerking, oral automatisms, involuntary movements, sudden behavioral arrest, tongue bite | CPS, Hypomotor, Automotor, Ictal Smiling, Behavioral Arrest, Aura | 4–40 | - |
| **FCD I + IIIa** | GTCS, behavioral arrest, sudden falls, eye deviation | GTCS, Behavioral Arrest | 4–34 | ~8% |
| **FCD IIb + IIIa** | Head nodding, smiling, limb automatisms, tongue bite, micturition | Ictal Smiling, Behavioral Arrest, Automatisms | 3–40 | ~5% |
| **FCD I + MTS** | GTCS, eye deviation, sudden falls | GTCS, Focal to Bilateral Tonic-Clonic Seizures | 4–58 | ~6–7% |
| **FCD I + II + MTS** | Multiple seizure types, resistant to AEDs | Mixed Seizure Semiology | 4–58 | ~15% |

**Supplementary table 3: Selection of molecular markers using bioinformatics tools data bases**

| **Gene** | **Associated Syndrome/Condition** | **Frequency/Expression** | **Reason for Selection** | **References** |
| --- | --- | --- | --- | --- |
| **MTOR** | FCD  RA  SKS | 0.9 (FCD)  0.6(RA)  0.9 (SKS) | mTOR pathway is a key player in cell growth and neuronal development. Dysregulation of mTOR has been linked to FCD. | Wong et al., 2015; Huang et al., 2017 |
| **DEPDC5** | Seizure  FCD | 0.9 (Seizure)  0.9 (FCD) | DEPDC5 is part of the mTOR signaling pathway and its mutations are associated with focal epilepsy and FCD. | Dibbens et al., 2013 |
| **RALA** | FCD | 0.75 | RALA, a small GTPase, is involved in signaling pathways that could affect neuronal development and synaptic plasticity, relevant to FCD. | Zhang et al., 2014 |
| **RHEB** | FCD | 0.65 | RHEB is involved in the regulation of the mTOR pathway, which plays a role in neuronal growth and synaptic function, making it a candidate gene for FCD. | Inoki et al., 2003; Pende et al., 2004 |
| **AKT3** | FCD | 0.98 | AKT3 mutations lead to abnormal cell growth and are involved in neurodevelopmental disorders that overlap with FCD features. | Kwiatkowski et al., 2014; Zhou et al., 2017 |
| **HTR6** | FCD | 0.67 | HTR6, a serotonin receptor, may influence neural network function and contribute to the pathophysiology of FCD-related seizures. | Vollenweider et al., 1997; Coller et al., 2013 |
| **PTEN** | FCD | 0.89 | PTEN mutations lead to neurodevelopmental disorders and are involved in the regulation of mTOR, thus playing a role in FCD pathophysiology. | Podsypanina et al., 1999; Li et al., 1997 |
| **KCNT1** | FCD  Seizure | 0.8 (FCD)  0.9 (Seizure) | KCNT1 mutations are linked to early-onset seizures and can contribute to neuronal hyperexcitability seen in FCD and epilepsy syndromes. | Tricarico et al., 2018 |

**Supplementary table 4: Scoring of IHC slide for FCD markers expressions**

| **Markers** | **AKT3** | **PTEN** | **MTOR** | **HTR6** | **RHEB** | **KCNT1** | **RALA** | **DEPDC5** |
| --- | --- | --- | --- | --- | --- | --- | --- | --- |
| FCD I | 28.99±1.20 | 15.90±0.76 | 103.63±4.12 | 16.50±0.68 | 10.60±0.78 | 03.01±0.09 | 04.02±0.03 | 02.01±0.01 |
| FCD II | 53.56±2.12 | 32.88±3.11 | 29.34±1.61 | 27.45±1.77 | 15.21±0.54 | 02.01±0.02 | 03.01±0.01 | 04.01±0.02 |
| FCD III | 121.21±3.21 | 53.03±2.51 | 109.97±5.76 | 46.29±2.33 | 18.060±0.25 | 04.01±0.01 | 03.01±0.02 | 03.01±0.01 |
